# TRIOPS: A deep learning framework for prediction of T cell receptor–MHC binding specificity

**DOI:** 10.64898/2026.06.30.735718

**Authors:** Nicholas R Rose, Cynthia M Ramirez, Lydia Mok, Christopher K Wong, Vanessa D Jonsson

**Affiliations:** Department of Biomolecular Engineering and Bioinformatics, University of California, Santa Cruz, California, USA; Department of Applied Mathematics, University of California, Santa Cruz, California, USA; Genomics Institute, University of California, Santa Cruz, California, USA

**Keywords:** T cell receptor, MHC restriction, HLA, deep learning, immune repertoire

## Abstract

T cell receptor (TCR) recognition is MHC-restricted, yet accurately predicting a TCR’s restricting HLA allele remains an open problem. We present TRIOPS, a dual-branch convolutional model with soft cross-attention that predicts TCR–MHC restriction from amino acid sequence alone. TRIOPS uses cross-reactivity–aware negative sampling by HLA pseudosequence similarity to reduce allele-boundary label noise, extending prediction to alleles absent from training. TRIOPS reaches a held-out AUC of 0.97 for paired TCRαβ and 0.92 for TCRβ-only inputs, generalizes to unseen receptors and HLA alleles, and after locus-specific calibration, assigns TCR clonotypes to their likeliest restricting allele across an individual’s HLA genotype. In TCGA tumors, TCR repertoires preferentially engage the expression-lost allele at HLA-A and HLA-B and the retained allele at HLA-C, recapitulating from bulk tumor RNA-seq the allele specific HLA loss previously linked to immune escape.

## Introduction

T cells recognize and eliminate infected or malignant cells by engaging antigens presented on major histocompatibility complex (MHC) proteins. Recognition is mediated by the T cell receptor (TCR), whose specificity arises from combinatorial V(D)J rearrangement. The resulting theoretical diversity, estimated at 10¹⁵–10²⁰ unique receptors^1,2^, enables broad immune surveillance but renders systematic experimental mapping of TCR–pMHC restriction intractable.

Central to this recognition is MHC restriction: TCRs engage with peptides only within the context of the MHC alleles that present them^3^, a constraint established during thymic selection. Positive selection acts on TCR engagement of self-peptide–MHC complexes in the thymus, producing a peripheral repertoire with intrinsic reactivity biased toward the selecting MHC alleles^4^. Resolving which HLA alleles predominantly restrict a given TCR collapses the antigen search to the peptides presentable by those alleles, narrowing the space of candidate TCR–antigen pairs for personalized cell therapies.

Class I and class II HLA are surveyed by CD8+ and CD4+ T cells respectively^5^, offering a mechanistic route to distinguish CD4 versus CD8 phenotype from sequence alone. Because restriction prediction requires no presented peptide, it extends to repertoire-scale data, where antigen identity is unknown and direct experimental pairing is intractable. Accurate TCR–HLA restriction prediction therefore addresses a layer of recognition upstream of, and presupposed by, peptide-level TCR–pMHC matching.

Computational models for predicting TCR–antigen restriction have advanced considerably but remain constrained in several ways. Peptide-dependent methods such as ERGO-I^6^, TCRAI^7^, NetTCR 2.2^8^ pMTnet^9^ and TAPIR^10^ require a known epitope, restricting their use to settings where the antigen is already defined, and independent benchmarks show that they generalize poorly to peptides underrepresented in training^11^.

Methods that predict TCR–MHC restriction without the peptide face their own limitations. CLAIRE^12^ is trained on experimentally validated pairs, but its per-allele performance is nonuniform. Many alleles near AUC 0.6 and few substantially above 0.8, and because it encodes HLA alleles as categorical name strings, it cannot predict alleles absent from training. Similarly, TAPIR^10^ predicts TCR–HLA pairs through peptide-masking augmentation, but was evaluated on only eight alleles, and its aggregate AUC of 0.81 obscures substantially weaker per-allele performance (e.g., 0.68 for HLA-A*03 despite abundant training data). For neither method are trained weights, model architecture, or source code publicly available.

DePTH2.0^13^ encodes both partners but is trained on statistically associated TCR β-chain (TCRβ)–HLA pairs identified by co-occurrence in population repertoires^14^ rather than on validated binding. The same population-level association is the basis for methods that impute an individual’s HLA genotype directly from their repertoire^15,16^. While a robust signal for that task, allele association is an indirect proxy for the physical restriction of any single receptor. TCRs are frequently cross-reactive across structurally related alleles rather than restricted to a single one^17^, and co-occurrence reflects HLA haplotype linkage as much as direct restriction. The resulting associations transfer poorly to experimentally validated pairs, a mismatch the DePTH2.0 authors have themselves documented^13^. DePTH2.0 is the only peptide-independent method with publicly available code and trained models.

A limitation shared across these methods is the generation of negative training data. TCR recognition is inherently degenerate: a single TCR can productively engage an estimated 10⁶ distinct pMHC ligands^18^, and this degeneracy extends to restriction itself, as structurally similar alleles present overlapping peptide repertoires and are recognized by overlapping sets of TCRs^19^. Random shuffling of TCR–MHC pairs that ignores allele similarity therefore introduces false negatives wherever cross-reactivity exists. For example, HLA-A*03:01 and HLA-A*11:01 differ at only four sequence residues in the peptide binding domain and share substantial TCR overlap across public datasets^20–24^, yet random shuffling labels a TCR binding either allele as a negative for the other. The resulting label noise corrupts the training signal and disproportionately penalizes discrimination at the allele boundaries where it matters most.

Here we present TRIOPS (T cell Receptor–Major Histocompatibility Complex Specificity), a deep learning model that predicts TCR–MHC compatibility directly from sequence. Rather than requiring a peptide, TRIOPS models MHC restriction itself, reflecting the MHC bias established during thymic selection. By representing MHC alleles as pseudosequences^25^, it utilizes shared features between related alleles to generalize beyond the alleles seen in training. This is essential given that the HLA system comprises thousands of alleles^26^, only a fraction of which have substantial representation in experimental datasets.

TRIOPS introduces three technical innovations: (1) cross-reactivity–aware negative sampling, in which negative TCR-MHC pairs are generated according to MHC sequence similarity, so that plausible associations between related alleles are preserved; (2) a dual-branch convolutional architecture trained with loss reweighting and contrastive regularization to sustain accuracy across alleles of widely varying representation; and (3) loci-specific False Positive Rate (FPR) calibration, which normalizes scores across MHC loci and enables each clonotype to be assigned to its most likely restricting allele across an individual’s full HLA genotype. We show consistent performance across HLA alleles and TCR chain configurations, generalization to unseen TCRs and HLA alleles, and improved accuracy over DePTH2.0. Applied to 8,906 TCGA pan-cancer samples, TRIOPS predictions recapitulate known clonal-expansion biology, recovering the expected dominance of oligoclonal CD8⁺ over polyclonal CD4⁺ T cell responses without supervision on T cell phenotype. At HLA-A and HLA-B, the allele most engaged by a patient’s TCR repertoire is preferentially the allele lost in tumors, whereas at HLA-C it is preferentially retained. This recovers allele-specific HLA loss, a known route of immune escape^27^, from the TCR repertoire alone.

## Results

### TRIOPS predicts TCR HLA restriction from receptor and allele sequence

TRIOPS predicts TCR–MHC restriction directly from amino acid sequence, producing both a per-pair compatibility score and a learned joint embedding. These support downstream tasks such as likelihood ranking of candidate TCR–HLA pairs and identifying predominantly targeted HLA alleles in patient samples (Fig. 1a). The model uses a dual-branch architecture that builds separate TCR and MHC representations before modeling their restriction (Fig. 1b). TCR sequences are encoded as the concatenated CDR1–CDR2–CDR3 of both TCR chains, with missing chains filled by placeholder tokens so that paired TCRαβ and single-chain inputs can be handled(Supplementary Fig. 1).

**Figure 1.**
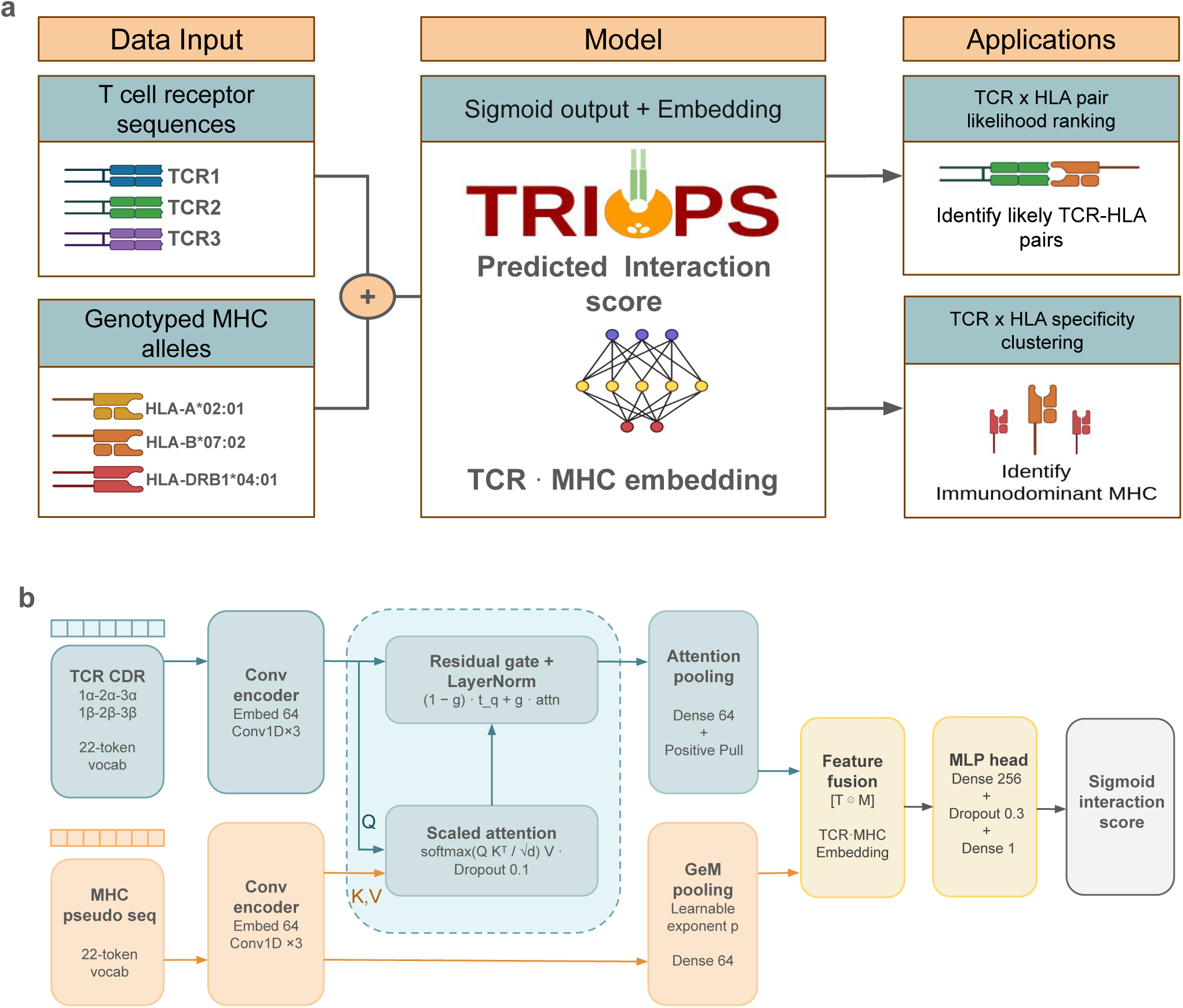
Overview and architecture of the TRIOPS framework for prediction of TCR–MHC specificity. a, Conceptual overview. T cell receptor sequences and genotyped MHC alleles are paired inputs to a model that predicts binding compatibility, returning an interaction score and a joint TCR·MHC embedding. Applications include TCR × HLA pair likelihood ranking and specificity clustering to identify immunodominant MHC. b, Neural network architecture. The six TCR CDR loops and the MHC pseudosequence are tokenized, embedded, and encoded through three Conv1D blocks each. In scaled cross-attention, the TCR supplies the query (Q) and the MHC the keys (K) and values (V), so each TCR position draws on its most relevant MHC features; a learned gate then merges this output with the original TCR representation. Attention pooling (TCR) and GeM pooling (MHC) yield 64-dimensional representations that are fused by element-wise multiplication and passed through an MLP head with sigmoid activation to produce the interaction score.

MHC alleles are represented by their NetMHCpan derived pseudosequences^25^. Within each branch, convolutional feature extractors learn local sequence motifs, and a cross-attention mechanism allows the TCR representation to attend over MHC features. The two branches are pooled and combined into a joint TCR·MHC embedding, which a multilayer perceptron (MLP) head maps to a sigmoid-activated compatibility score (Fig. 1b). A PositivePull regularizer draws together the embeddings of true restriction pairs, counteracting the fragmentation that sequence sparsity and class imbalance would otherwise induce.

Negatives are sampled to respect cross-reactivity: only mismatched alleles at a pseudosequence Levenshtein distance (d) of at least 10 amino acids from a TCR’s true restricting allele are used as negatives, while closer alleles, which may be genuinely cross-reactive, are excluded (Supplementary Fig. 2). Among allele pairs sharing paired TCRs, normalized overlap persists across short-to-intermediate distances but falls to near zero at d ≥ 10 (Supplementary Fig. 2d), marking the smallest distance at which a mismatched allele can be treated as a true negative. This boundary coincides with the leveling of the cluster separation ratio at d = 10–12, where intracluster pairs remain substantially more similar than intercluster pairs (Supplementary Fig. 2b,c)

#### Accurate restriction prediction across HLA class I and II and TCR chain configurations

We evaluated TRIOPS by six-fold cross-validation, and reported ensemble predictions (mean interaction score across folds) for all metrics. On the held-out evaluation set, paired-input AUCs ranged from 0.82 to 1.00 and most alleles above 0.89 (Fig. 2a). Class II alleles performed as strongly as class I (per-allele AUC 0.89–1.00 versus 0.82–0.99; Fig. 2a) despite the marked training-set imbalance in favor of class I (Supplementary Fig. 3).

**Figure 2.**
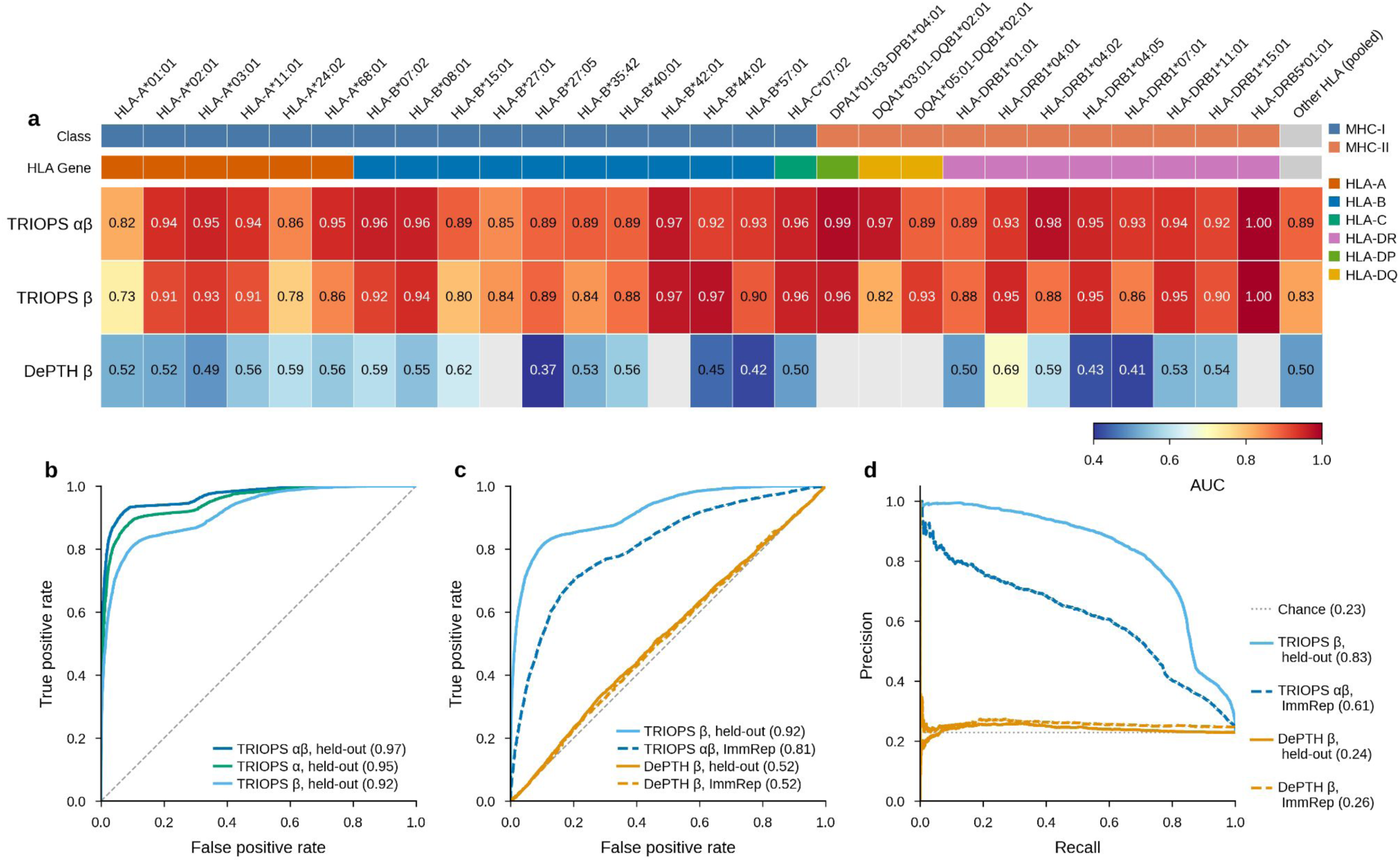
Predictive performance of TRIOPS and comparison with DePTH2.0. a, Per-allele AUC heatmap for TRIOPS (paired TCRαβ and TCRβ-only) and DePTH2.0 (TCRβ-only). Alleles ordered by gene group; color strips indicate MHC class and gene. Grey cells: alleles not evaluable by DePTH2.0. b, ROC curves by chain configuration (paired, AUC = 0.97; α-only, 0.95; β-only, 0.92). c, Head-to-head ROC. Solid: held-out (TRIOPS β, 0.92; DePTH β, 0.52). Dashed: IMMREP (TRIOPS αβ, 0.81; DePTH β, 0.52). d, Precision-recall curves for the same comparisons.

Aggregated across alleles, paired TCRαβ inputs achieved the highest pooled ROC-AUC (0.97), followed by TCRα-only (0.95) and TCRβ-only (0.92) (Fig. 2b). Both single-chain configurations were strongly predictive, consistent with each chain carrying information relevant to restriction.

We then evaluated TRIOPS on the IMMREP 2023^28^ TCRαβ benchmark, restricted to TCRs absent from every training fold and the held-out evaluation dataset. On these entirely unseen receptors, paired αβ inputs reached a ROC-AUC of 0.81 (Fig. 2c), with a TCRαβ precision–recall AUC of 0.61 (Fig. 2d). Performance remains well above chance (precision–recall AUC of 0.24), indicating that TRIOPS captures generalizable TCR–MHC sequence determinants rather than memorizing training receptors.

Because TRIOPS encodes each allele by its pseudosequence, we assessed whether predictions extend to alleles absent from training. Fifteen pseudosequences, corresponding to 16 HLA alleles, were withheld across all six folds while their associated TCRs were retained, thereby isolating MHC novelty from receptor novelty (n = 56,703 pairs, 9.3% binders; Supplementary Fig. 4). The six-fold ensemble achieved a ROC-AUC of 0.92 on this holdout. Per-allele performance ranged from 0.43 (HLA-A*25:01) to 1.00 (HLA-A*02:56 and HLA-C*04:01), and the allele-balanced macro-average AUC was 0.87. The macro-average provides the more representative summary, as a single allele (HLA-B*07:02) constituted 83% of holdout pairs.

#### TRIOPS outperforms DePTH2.0 on validated pairs and unseen receptors

The public implementation of DePTH2.0 accepts only TCRβ, and a fixed set of 85 class I and 250 class II alleles, whereas TRIOPS represents each allele by its pseudosequence and so extends to alleles not encountered during training. Across the alleles that DePTH2.0 accepts, TRIOPS exceeds DePTH2.0 AUC at every HLA allele (TRIOPS 0.82–1.00 for paired TCRαβ and 0.73–1.00 for TCRβ-only; DePTH2.0 0.37–0.69; Fig. 2a); DePTH2.0 values near 0.50 reflect its evaluation outside the co-occurrence regime on which it was trained. TRIOPS’ advantage over DePTH2.0 was similar on the IMMREP benchmark of unseen TCRs^28^, where DePTH2.0 received a precision-recall AUC of 0.24 for TCRβ (Fig. 2d), indicating that DePTH2.0 retains little predictive signal on receptors beyond its training distribution.

This disparity is consistent with the two models learning distinct signals. A body of work infers HLA genotype from bulk TCR repertoires by exploiting the population-level association between public clonotypes and HLA background^15,16^, and DePTH2.0 operationalizes this co-occurrence signal for prediction. TRIOPS instead learns from experimentally verified restriction pairs. Co-occurrence reflects shared immune exposure and linkage between public clonotypes and an individual’s HLA background rather than the physical compatibility of a receptor with an allele, and a model fitted to the former does not necessarily transfer to per-pair restriction. CLAIRE, the closest peptide-independent comparator, releases neither trained weights nor a packaged model, precluding a direct benchmark.

#### Learned representations recover HLA allelic structure and TCR-MHC contact domains

TCR–HLA joint embeddings from positive validation pairs are separated by HLA gene group: On the 23 alleles with at least 20 representatives, embedding coherence exceeded the permutation null for every metric: k-NN purity 0.989 (null 0.284), adjusted Rand index (ARI) 0.224, normalized mutual information (NMI) 0.639 and silhouette coefficient 0.357 (Fig. 3abc). The ratio of between-cluster to within-cluster distance was 2.62 at the allele level (95% CI 2.568 to 2.667) and 1.83 at the class level (95% CI 1.796 to 1.861), both exceeding the no-separation value of 1.0 (Fig. 3d). Cosine-distance distributions increased with allelic and class separation: within-allele (median 0.016, IQR 0.008-0.030), between-allele within-class (median 0.049, IQR 0.033-0.076), and between-class (median 0.081, IQR 0.063-0.111) (Fig. 3e).

**Figure 3.**
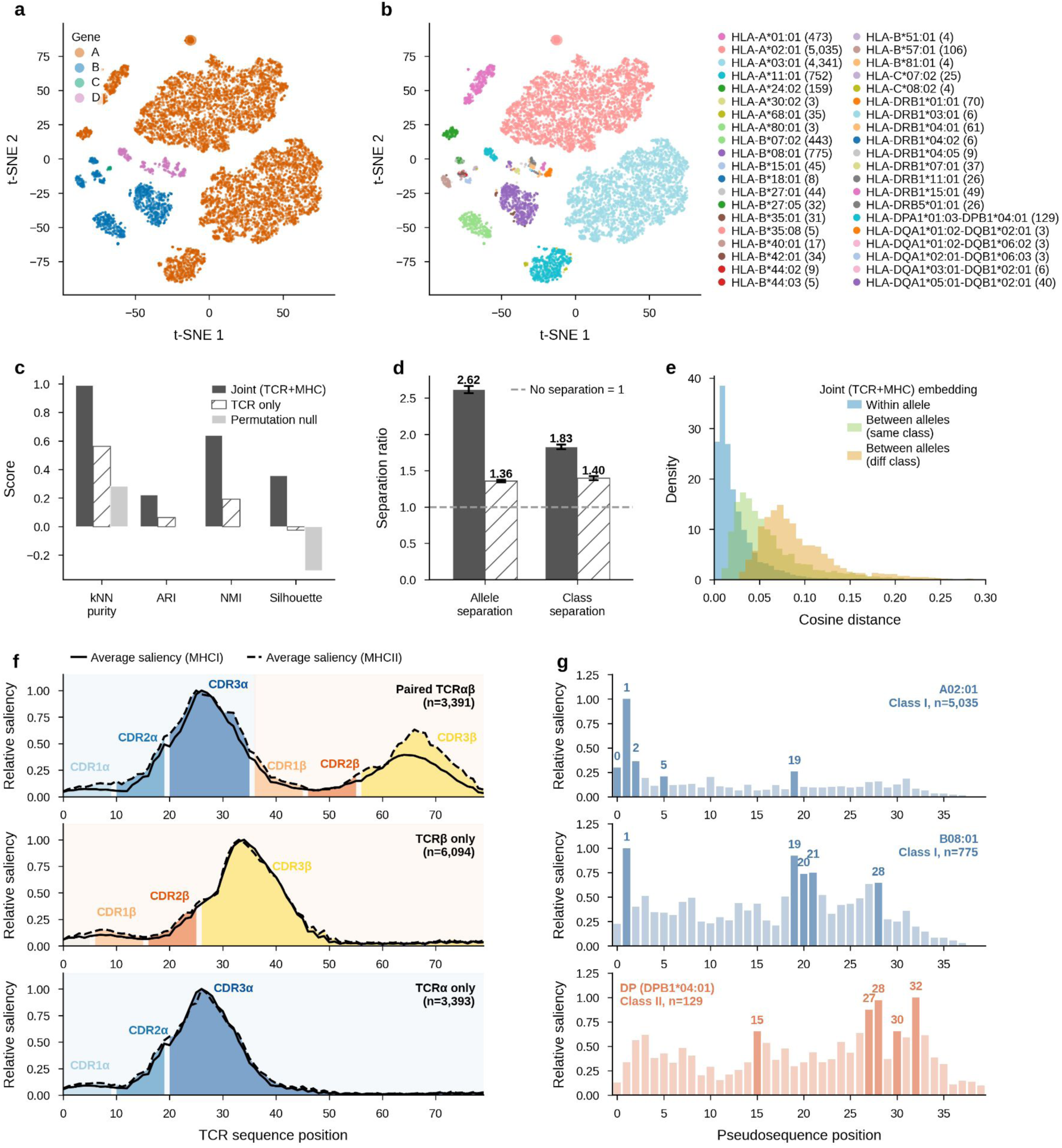
TRIOPS embedding structure, coherence analysis, and positional saliency. All panels use the fold-4 model. a, t-SNE of fusion embeddings colored by HLA gene group. b, Same projection colored by individual allele (top 40 by frequency). c, Coherence metrics (kNN purity, ARI, NMI, silhouette) for the joint TCR·MHC embedding and a TCR-only embedding, each compared against the permutation null. d, Separation ratios (between-over within-group cosine distance) at the allele and class level for the joint embedding (2.62 and 1.83) and the TCR-only embedding (1.36 and 1.40); dashed line marks no separation (ratio = 1). e, TCR·MHC cosine distance distributions for within-allele, between-allele same-class, and between-class pairs. f, TCR positional saliency stratified by MHC class. Top, paired (n = 3,391); middle, β-only (n = 6,094); bottom, α-only (n = 3,393). g, MHC pseudosequence saliency for A*02:01 (n = 5,035), B*08:01 (n = 775), and DPA1*01:03-DPB1*04:01 (n = 129).

Positional saliency analysis on the TCR input data showed that, for paired inputs, CDR3α and CDR3β carried the highest average gradients, followed by CDR2α and CDR2β (Fig. 3f). Saliency for TCRβ-only inputs concentrated in CDR3β and CDR2β, and for TCRα-only inputs in CDR3α and CDR2α. MHC pseudosequence saliency was allele-specific (Fig. 3g). Both class I alleles showed a dominant peak at pseudosequence position 1 but differed in their secondary positions: HLA-A*02:01 at positions 2 and 19, and HLA-B*08:01 at positions 19–21 and 28. The class II allele HLA-DPA1*01:03-DPB1*04:01 demonstrated high saliency across both halves of the pseudosequence, with peaks in the N-terminal DPA1-derived region (position 15) and the C-terminal DPB1-derived region (positions 27–32). Saliency localized to the complementarity-determining loops, with contributions from both the somatically recombined CDR3 and the germline-encoded CDR2 that contacts the MHC α-helices. This is consistent with a model that integrates both sources of sequence variation in predicting restriction.

#### TRIOPS predictions recapitulate Class I restricted clonal dominance across TCGA tumors

We applied TRIOPS to a TCRβ repertoire dataset reconstructed from 8,906 TCGA pan-cancer tumor RNA-seq samples^29^. HLA genotypes were obtained from the TCGA pan-cancer consensus HLA typing benchmark dataset^30^ and resolved to a maximum of two alleles per locus at two-field resolution. HLA allele-specific expression was quantified using arcasHLA-quant^31^ on chromosome 6 reads extracted from TCGA tumor RNA-seq BAM files. Each TCRβ clonotype was scored using TRIOPS against every HLA allele carried by the patient. Raw scores were calibrated to a locus-specific 20% FPR threshold mean before comparison (Methods, Supplementary Fig. 5), and each clonotype was assigned to the allele with the largest calibrated margin.

TRIOPS produced a 192-dimensional joint TCR·MHC embedding for each of 417,400 argmax-assigned TCR–HLA pairs across 8,906 patients; visualized by t-SNE (50,000-pair proportional subsample, perplexity 30; Methods), the embeddings separated by MHC class (Fig. 4a) and resolved into allele-specific clusters, the largest being HLA-A*02:01 (79,328 pairs) (Fig. 4b). Within-patient locus fraction by HLA locus across n = 7,515 patients; HLA-A contributes the largest single-locus share (median ≈ 0.37), exceeding every other locus (next highest HLA-DR, 0.15; HLA-C 0.10, HLA-B 0.09, HLA-DQ 0.09, HLA-DP 0.07) (Fig. 4c). Predicted class I clonotypes were also retained at higher confidence margins than predicted class II (median class-level margin 0.115 vs 0.089; allele-level 0.098 vs 0.032), the gap widening at the allele level where the class II margin was lowest (Fig. 4d).

**Figure 4.**
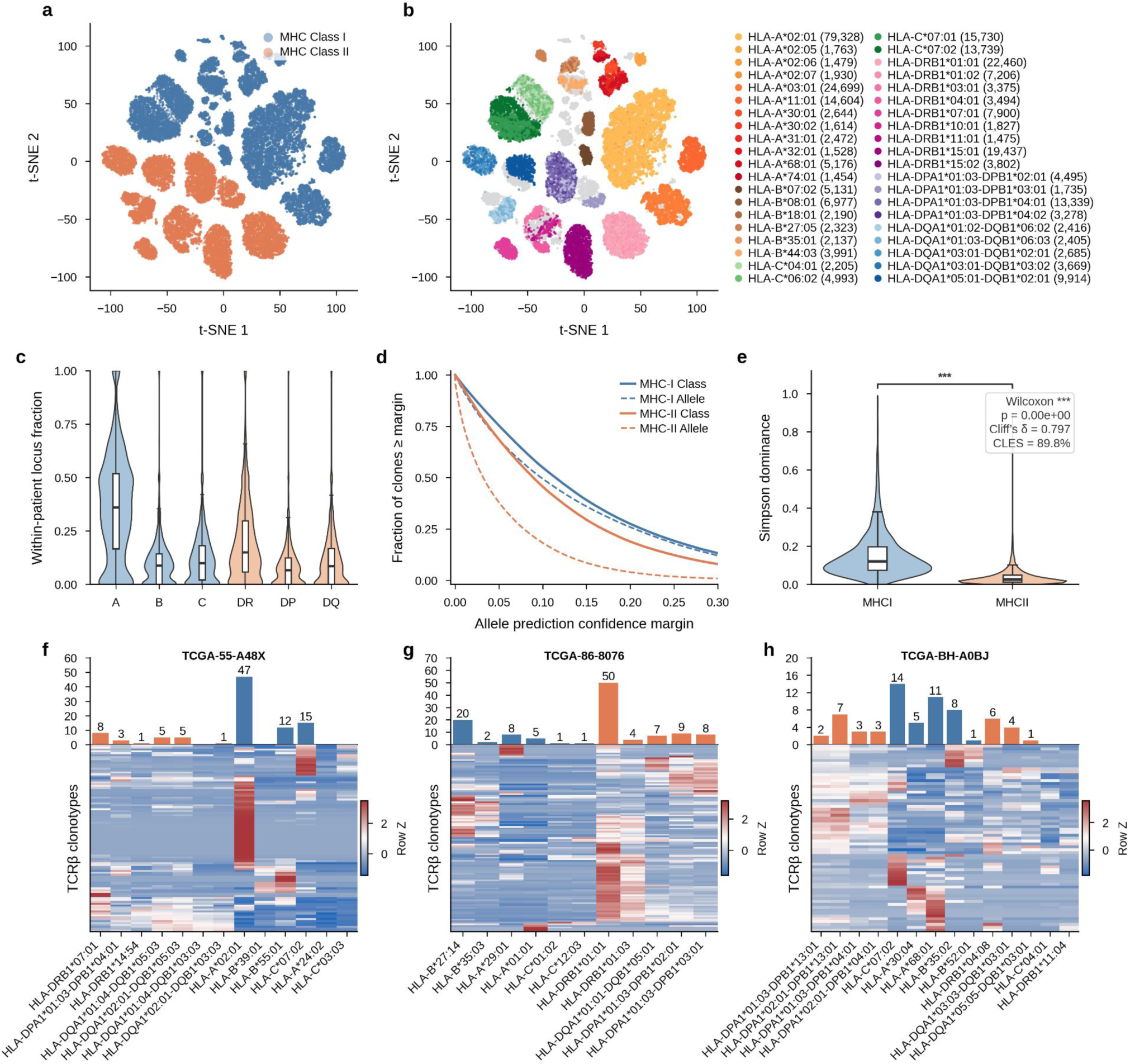
TRIOPS predictions in TCGA reveal patient-specific immune specificity landscapes and recapitulate T cell biology. a, t-SNE of argmax TCRβ–HLA assignments (n ≈ 417,400 clonotype assignments across 8,906 patients) colored by MHC class. b, Same projection colored by allele, with gene-family shading. c, Within-patient locus fraction by HLA locus across n = 7,515 patients; HLA-A contributes the largest single-locus share (median ≈ 0.37). d, Fraction of assigned clonotypes whose prediction confidence margin exceeds a given threshold, for MHC-I and MHC-II assignments. Class curves (solid) give the margin between the top-ranked allele and the best allele of the opposing class; allele curves (dashed) give the margin between the top two alleles within the assigned class. Class I assignments retain higher margins than class II at both levels. e, Within-class Simpson dominance for class I– versus class II–targeting TCRβ clonotypes (Wilcoxon P < 2.2 × 10⁻¹⁶; Cliff’s δ = 0.797; CLES = 89.8%). f, FPR-calibrated TRIOPS score heatmap (rows z-scored, hierarchically clustered) with allele-level clonotype counts above, for TCGA-55-A48X, a predominantly class I–restricted repertoire. g, Same for TCGA-86-8076, a predominantly class II–restricted repertoire. h, Same for TCGA-BH-A0BJ, spanning both classes.

Within-class Simpson dominance was higher for predicted class I– than for predicted class II TCRβ clonotypes (Cliff’s δ = 0.797; Wilcoxon P < 2.2 × 10⁻¹⁶; n = 8,906 patients, CLES = 89.8%; Fig. 4e). This recapitulates the established observation that CD8⁺ T cell responses in tumors are dominated by a few highly expanded clonotypes, whereas CD4⁺ repertoires retain greater clonal diversity^32^. Because TRIOPS is supervised only on restriction, not on clonality or lineage, this CD8-versus-CD4 expansion asymmetry emerging from its class I/II predictions indicates that predicted MHC-class targeting carries lineage-linked clonal structure the model was never trained on. As a further independent test in lung adenocarcinoma (LUAD), TRIOPS-predicted MHC-targeting clone counts correlated with CIBERSORT-estimated^33^ T cell proportions, demonstrating concordance between two orthogonal features of the same sample. (Supplementary Fig. 6).

At the single-patient level, each TCRβ clonotype is assigned to the carried allele giving the highest calibrated margin, distributing a patient’s repertoire across their HLA alleles (Fig. 4f–h). The resulting profiles vary widely: TCGA-55-A48X is dominated by class I–targeting clonotypes (Fig. 4f) and TCGA-86-8076 by class II–targeting clonotypes (Fig. 4g), whereas TCGA-BH-A0BJ spans both (Fig. 4h).

#### TCR engagement is associated with allele specific expression loss of HLA-A and HLA-B and retention of HLA-C across cancers

Using TRIOPS per-clonotype restriction predictions for the TCGA pan-cancer cohort, we quantified each allele’s TCR engagement as the read-weighted abundance of the clonotypes assigned, where the engagement allele of a heterozygous pair is the one with the greater TCR engagement. Allelic expression is measured by allele-resolved RNA-seq B-allele frequency (BAF). Because BAF is measured from RNA, this readout captures allele-specific loss at the level of expression, encompassing both genomic loss and transcriptional silencing. Twenty three cancer types were analyzable.

Across cancers, we summarized each cancer type as a concordance probability c: the covariate-adjusted probability, in a typical cancer type, that the engaged allele is the lost one (null 0.5; Firth regression adjusting for tumor purity and read depth, aggregated across cancers; Methods). The class I loci separated cleanly, the effect was strongest at HLA-B (c = 0.65; 23/23 cancers lost-engaged), unanimous but weaker and more dispersed at HLA-A (c = 0.61; 23/23), and reversed at HLA-C (c = 0.37; 0/23), where the engaged allele is preferentially retained (Fig. 5a). Because the sign test counts only direction, its P value saturates at the unanimity floor for 23 cancers (2 × (1/2)²³ = 2.4 × 10⁻⁷); we therefore report directional counts throughout and assess effect magnitude with per-cancer tests.

**Figure 5.**
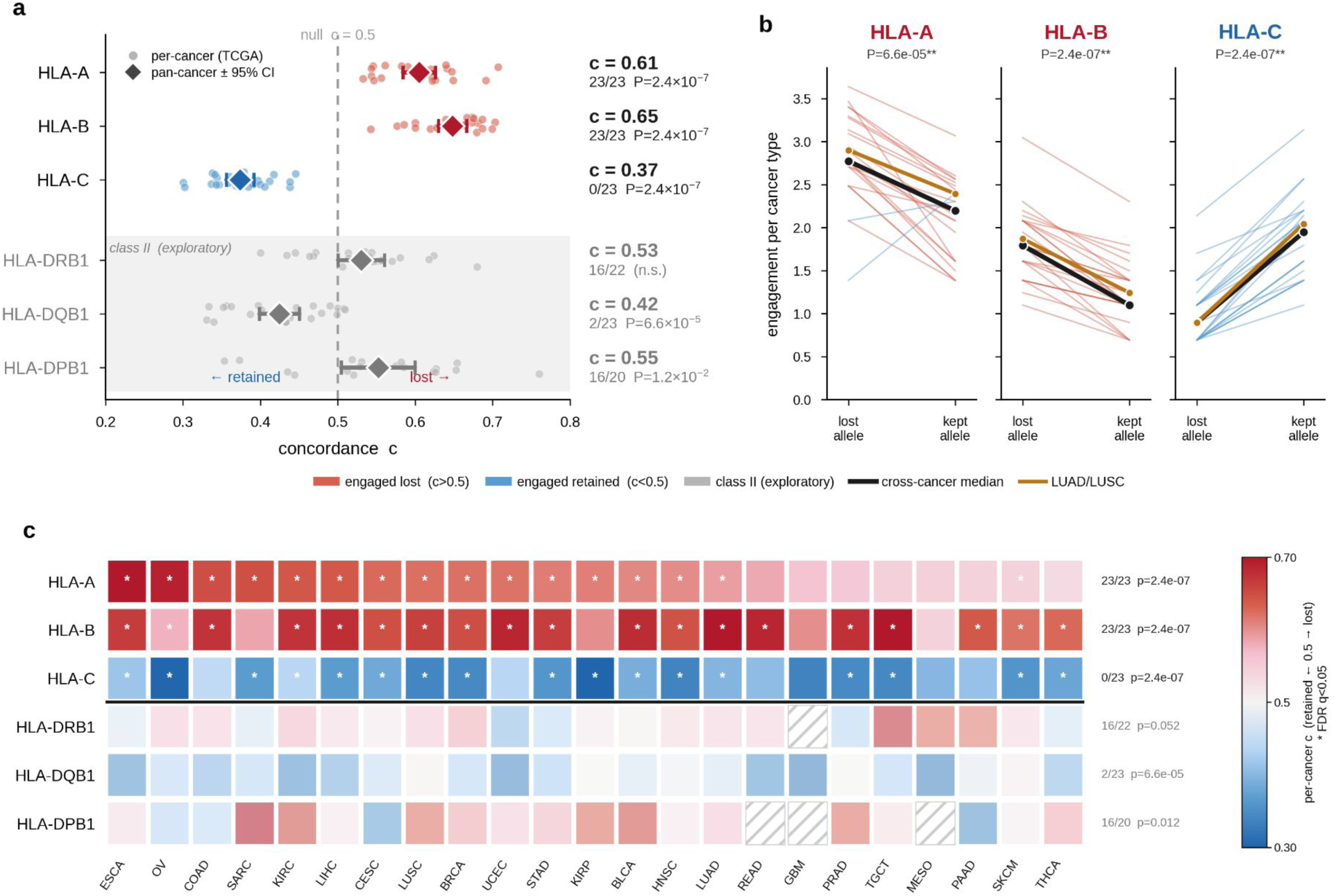
TCR engagement is associated with allele-specific HLA loss across cancers. a, Covariate-adjusted concordance c by HLA locus, where c is the probability that the more-engaged allele is the under-expressed (lost) member of a heterozygous pair (null c = 0.5; purity- and depth-adjusted Firth logistic regression). Diamonds, pan-cancer estimate with 95% CI; points, individual cancer types; red, engaged allele preferentially lost (c > 0.5); blue, preferentially retained (c < 0.5). Class II loci (gray) are exploratory. At right, the number of cancer types in the lost-engaged direction over the number testable, and P (two-sided exact binomial sign test across cancer types). b, Within-tumor paired engagement at the lost (BAF < 0.5) versus retained allele for the three class I loci. Each thin line is one cancer type’s median engagement; black, cross-cancer median; gold, LUAD/LUSC. Downward slopes at HLA-A and HLA-B indicate higher engagement at the lost allele; the upward slope at HLA-C indicates higher engagement at the retained allele. P, two-sided exact binomial sign test across cancer types. c, Per-cancer concordance c for each locus × cancer type, estimated without pooling; class I loci above the rule, class II below. Color encodes direction and magnitude (red, engaged-lost; blue, engaged-retained; intensity, |c − 0.5|; range 0.30 to 0.70); asterisks mark cells significant at q < 0.05 (Benjamini–Hochberg across all finite locus × cancer cells). At right, cancer-type count in the consistent direction over the number testable, and sign-test P.

Within individual tumors, we compared TCRβ engagement at the lost (BAF < 0.5) versus retained allele of each informative pair, defining the lost allele by expression alone (Fig. 5b). Engagement was higher at the lost allele in 21/23 (HLA-A) and 23/23 (HLA-B) cancers, and in 0/23 at HLA-C — a reversal that holds each pair’s tumor context fixed, independent of the across-cancer aggregation.

Resolved by individual cancer type (Fig. 5c), the effect was broadly distributed rather than concentrated in a few cancers: per-cancer estimates reached q < 0.05 (Benjamini–Hochberg) in 23/23 (HLA-A) and 23/23 (HLA-B) cancers and reversed at HLA-C.

Engagement and BAF both scale with allelic expression, so a uniform engagement–loss association could in principle be an expression artifact rather than immune selection. The reversal at HLA-C, where the engaged allele is instead retained, therefore points to locus-specific biology rather than a measurement artifact: CD8⁺ immunoediting at HLA-A and HLA-B, NK missing-self constraints at HLA-C. This locus-dependent pattern is consistent with allele-specific HLA loss previously linked to immune escape^27^, recovered here from expression rather than genomic copy number, and with engaged alleles marking the targets of immune selection.

Class II predictions showed no clear pattern and are treated as exploratory: HLA-DRB1 and HLA-DPB1 leaned weakly toward loss (c = 0.53, 0.55), while HLA-DQB1 reversed toward retention (c = 0.42). Lacking the locus-consistent reversal that excludes expression covariation for class I, we do not interpret the DQB1 signal.

## Discussion

We presented TRIOPS, a deep learning model that predicts TCR–MHC restriction directly from sequence, without a peptide. Its central methodological choice is cross-reactivity–aware negative sampling: rather than shuffling TCR–MHC pairs at random—which mislabels structurally related, cross-reactive alleles as negatives—TRIOPS draws negatives by MHC pseudosequence similarity and learns graded relationships among related alleles rather than hard allele boundaries. A single architecture handles α-only, β-only and paired αβ receptors, sustains accuracy for sparsely represented alleles through loss reweighting and contrastive regularization, and calibrates scores across loci so that every clonotype can be assigned to its most likely restricting allele across a patient’s full HLA genotype.

On experimentally validated binding data, TRIOPS outperformed DePTH2.0 and generalized to unseen receptors, a gap consistent with DePTH2.0 being trained on population-level allele co-occurrence rather than validated binding^13^. A direct comparison against CLAIRE^12^, the closest peptide-independent predictor, was not possible: it releases neither trained weights nor model code and encodes alleles as categorical names rather than sequence, precluding evaluation on the unseen alleles and pseudosequence-resolved pairs assessed here. TCRα-only inputs outperformed TCRβ-only inputs (AUC 0.95 versus 0.92), consistent with crystallographic evidence that the α-chain contributes dominant contacts to the MHC α-helices^34^. That this is recovered from sequence alone, with saliency localizing each single-chain model’s signal to its own CDR loops, indicates genuine α-chain restriction features rather than dataset asymmetry.

Applied to TCGA pan-cancer TCR repertoires, TRIOPS reflects the repertoire it receives, not the repertoire in vivo; its outputs are predictions conditioned on the recovered input rather than direct biological readouts. The predicted landscape is dominated by HLA-A (e.g., HLA-A*02:01, 79,328 clonotypes; HLA-B*08:01, 6,977), which we read as a detection-floor artifact of untargeted assembly rather than a true difference in response magnitude: TCGA RNA-seq is unenriched for TCRs (locus coverage ≈0.04×), TRUST4 recovery scales with locus expression (Spearman r ≈ 0.35)^35^, and reliable detection is confined to the ∼0.1–1% repertoire-frequency range. TCR recovery from TCGA samples by TRUST4 is therefore biased toward clones that occupy a large fraction of their host repertoire, regardless of restricting locus. Dedicated TCR-enriched or single-cell repertoire sequencing can recover the lower-frequency clones that unenriched TCGA RNA-seq misses, mitigating this bias in future cohorts.

Allele-specific HLA loss is an established mechanism of immune escape in cancer, and at the DNA level it is widespread: across a pan-cancer cohort of 83,644 samples, Montesion et al.^36^ found somatic HLA-I loss of heterozygosity in 17% of tumors, establishing it as a common route to immune evasion. That study also reported a nonlinear relationship between HLA-I LOH and tumor mutational burden, with loss most frequent at intermediate burden and declining above 30 mutations per megabase, indicating that the mechanism is favored in some tumor contexts more than others. Using LOHHLA on matched tumor-normal exomes, McGranahan et al.^27^ reported enrichment of subclonal neoantigens in NSCLC predicted to bind to the lost allele, evidence that allele-specific loss is shaped by immune targeting.

We recover the same directional relationship from the TCR repertoire side, substituting the TCRs a patient is predicted to engage for the antigens that patient is predicted to present: the allele a patient’s TCRβ repertoire most engages is preferentially the down-expressed one at HLA-A and HLA-B (concordance c = 0.61 and 0.65) and the retained one at HLA-C (c = 0.37). The two analyses converge from opposite measurements, and ours requires neither a neoantigen prediction nor a matched exome. Because TRIOPS never saw allelic expression, HLA genotype, or loss during training, the direction is imposed by neither the model nor the analysis. The cross-cancer heterogeneity in Fig. 5c is consistent with this context dependence, the per-cancer concordance varying in magnitude across tumor types while preserving the locus-level direction.

We frame this convergence with a caveat: TRUST4 reconstructs the bulk intratumoral TCRβ repertoire and TRIOPS assigns each clonotype to its restricting allele, not to a target antigen; TCRβ engagement mass therefore measures how much of a patient’s recoverable repertoire an allele restricts, not how much tumor-directed pressure it bears. Because intratumoral repertoires contain substantial bystander clones, our engagement–loss concordance is an association consistent with immunoediting^37^ rather than a demonstration of it: McGranahan et al.^27^ bridged to tumor reactivity through neoantigen binding, a step our restriction-based measure omits.

The three class I loci are unequally exposed to this caveat. HLA-C is the most robust to the restriction versus reactivity caveat: its retention (c = 0.37; engaged allele lost in 0 of 23 cancers) is most parsimoniously explained by an NK-cell “missing-self” constraint. HLA-C is the dominant ligand for inhibitory killer-cell immunoglobulin-like receptors and dominant educator of the NK compartment; its constitutively low surface expression marks a locus whose primary role is NK tuning rather than antigen presentation^38^. Independent repertoire analyses corroborate this from the TCR side: DeWitt et al.^15^ reported weaker statistical association between TCRs and HLA-C than other class I loci and attributed it to the same low surface expression and KIR-mediated NK interaction.

NK cell responsiveness is itself calibrated by the engagement of inhibitory receptors with self class I MHC, a process termed licensing or education, so that NK cells tuned to a given HLA-C allele become competent to respond when that allele is subsequently lost^39,40^. Losing HLA-C therefore exposes a cell to NK missing-self lysis irrespective of whether its engaging T cell clones are tumor-reactive, so retention is expected whether the clones are effectors or bystanders, consistent with tumors commonly undergoing allelic HLA loss but rarely complete class I loss^41^. Our contribution is the observation that the endogenously most-engaged HLA-C allele is preferentially retained. The reported loss of HLA-C*08:02 under a dominant KRAS-G12D-restricted clone in adoptive transfer^42^ fits rather than opposes this view: it reflects a focused therapeutic response at a single allele, whereas our estimand is the net effect of the endogenous repertoire across patients, where HLA-C is retained.

HLA-A and HLA-B show the converse: at both, the engaged allele is preferentially the lost one. The association is strongest and most reproducible at HLA-B, where the engaged allele is the lost one in 23 of 23 cancers in the per-cancer concordance analysis and carries higher engagement than the retained allele in 23 of 23 cancers within tumors, consistent with active immunoediting. HLA-B is independently the locus most clearly tied to immunotherapy outcome. HLA-B alleles present roughly twice as many self-peptides as HLA-A (∼430 versus ∼212)^43^ and sit toward the “generalist” end of a promiscuity–expression trade-off^44^. In 1,535 checkpoint-blockade-treated patients, Chowell et al.^45^ found that the HLA-B44 supertype conferred extended survival whereas the HLA-B62 supertype or somatic HLA-I loss of heterozygosity predicted poor outcome, with maximal heterozygosity across HLA-A, HLA-B and HLA-C improving overall survival. That the locus bearing our strongest engagement–loss signal is also the one where germline restriction and its somatic loss most influence survival links our molecular result to a clinical phenotype, though we do not test the survival relationship directly here. HLA-A is directionally identical but weaker (c = 0.61; engaged allele lost in 23 of 23 cancers, higher engagement at the lost allele in 21 of 23 within tumors). Because restriction is not reactivity, we state both as engagement tracking loss consistent with active immunoediting rather than demonstrated CD8-driven selection; the HLA-A signal is the more exposed to the caveat that the engaged clones may be bystanders rather than tumor-reactive T cells.

This repertoire-side result mirrors what genomic analyses recover from the antigen-presentation side: across 6,319 primary and metastatic tumors, Martínez-Jiménez et al.^46^ found that focal HLA-I loss of heterozygosity preferentially eliminates the allele presenting the largest predicted neoepitope repertoire, evidence that the lost allele is the immunologically active one. The convergence is notable because the two measurements are orthogonal, neoepitope presentation on one side and TCR engagement on the other, yet both single out the lost allele. The same publication also found genetic immune escape prevalence to be generally consistent between primary and metastatic tumors, which supports our reading that treatment-naive primary tumors still capture editing: because TCGA samples are predominantly treatment-naive primary tumors and allele-specific HLA LOH is largely subclonal^27^, primary-tumor RNA captures an earlier, finer-grained expression phase of editing that precedes the coarser copy-number endpoint.

The detection asymmetry has a structural basis that reinforces the detection floor interpretation of the dominance landscape. The greater promiscuity of HLA-B alleles distributes a comparable response across more epitopes and pushes more HLA-B clones below TRUST4’s floor. The same dilution applies across the class I/II boundary, where class II clonotypes show lower per-allele dominance than class I (Cliff’s δ = 0.797). The asymmetry is thus a predicted consequence of sampling a broad repertoire through a shallow assay—not evidence that HLA-B- or class II-restricted responses are weaker.

Several limitations bound these conclusions. The engagement measure indexes restriction, not antigen reactivity, so the immunoediting reading is associational; class II results are exploratory, with no consistent directional concordance (HLA-DRB1 c = 0.53, HLA-DPB1 c = 0.55; HLA-DQB1 c = 0.42). The cross-sectional TCGA design captures a single point in tumor evolution and immune response, so the dynamic interpretation of active immunoediting at HLA-A and HLA-B is inferred from one snapshot rather than observed across time.

More broadly, any immune-infiltrated tissue with bulk RNA-seq is, in principle, a substrate for resolving the HLA-restricted clonal landscape underlying anti-tumor T cell responses^47^. TRIOPS supports two clinical modes: pairing a patient’s full repertoire against their genotype to assign each clonotype to its restricting allele and identify dominant restricting alleles, information inaccessible to bulk deconvolution^33^; or, given a neoantigen predicted to bind a specific allele, scoring every repertoire TCR against that allele as a pre-filter for TCR-T discovery. The latter addresses a concrete bottleneck: ∼80% of TCR-T trials are restricted to HLA-A*02^48^, and extending restriction prediction across diverse alleles—even at reduced accuracy on unseen TCRs (AUC 0.81 paired/0.77 beta only)—could broaden eligibility and reduce disparities in access to TCR-based therapeutics.

## Methods

### Data collection and harmonization

Experimentally validated TCR–pMHC binding data were downloaded from TCR3d^20^, VDJdb^21^, IEDB^22^, McPAS-TCR^23^, and PATCRdb^24^ (accessed 2024). Records were filtered to human TCRs with confirmed binding to human HLA alleles (HLA-A, -B, -C, -DR, -DP, -DQ). All sequences were harmonized to IMGT nomenclature^49^. CDR1 and CDR2 sequences were extracted from IMGT-gapped V-gene reference sequences and extended with flanking framework residues to 10-amino-acid fixed-length representations. CDR3 sequences were extracted according to IMGT definitions, truncated at the conserved F/W of the F-G-X-G motif. CDR loops were concatenated with hyphen delimiters (TCRα = CDR1α-CDR2α-CDR3α; TCRβ = CDR1β-CDR2β-CDR3β). Missing chains were encoded as “X-X-X”. MHC alleles were represented using 34-residue NetMHCpan^25^ pseudosequences (Supplementary Fig. 1). After deduplication, the positive dataset comprised 136,428 unique TCR–HLA pairs.

### HLA pseudosequence similarity and shared TCR usage

Pairwise Levenshtein distances were computed and alleles grouped by average-linkage hierarchical clustering. Group coherence was assessed across cut heights from 4 to 12 using a separation ratio R(d), the median intercluster distance divided by the median intracluster distance (95% confidence intervals from 200 bootstrap replicates). Shared TCR usage was measured between HLA alleles in our dataset, for the α chain, β chain, or both chains jointly. For every allele pair sharing at least one identical TCR, a normalized overlap was computed as the number of shared TCRs divided by the summed sizes of the two TCR sets, averaged across sharing pairs at each integer levenshtein distance.

### Negative data generation

Negative examples were generated by permuting the allele while retaining the TCR sequence. Candidates were rejected if the Levenshtein distance between original and replacement pseudosequences was ≤9 (retaining only distance ≥10). Sampling was stratified to approximately 65% cross-gene and 35% within-gene negatives. A top-off procedure ensured a ≥2:1 negative-to-positive ratio per allele, with the Levenshtein distance threshold relaxed to ≥9. The final dataset comprised 578,474 pairs (positive-to-negative ratio ∼1:3.6).

### Data augmentation and partitioning

Paired TCRαβ entries were augmented by chain masking (α-only and β-only versions), applied independently within each partition to prevent leakage. Data were stratified by allele, label, and chain status into a fixed held-out set, with remaining data divided into six cross-validation folds (∼80/20 train/validation). Held-out pairs were not present in training, though individual TCR sequences could appear in training paired with different MHC alleles; generalization to entirely unseen TCRs is evaluated separately on the IMMREP 2023 benchmark.

### Model architecture

TCR and MHC sequences were encoded using a 22-character vocabulary and embedded into 64-dimensional representations. Each branch processes sequences through three convolutional blocks (kernel size 3, filters 64→128→256, batch normalization, max pooling after blocks 1–2). Cross-attention projects TCR features to queries and MHC features to keys/values via 128-filter convolutions; scaled dot-product attention with dropout (p = 0.1) is combined with the original TCR features through a learned residual gate (init 0.2) and layer normalization. Attention pooling aggregates TCR features; GeM pooling (trainable p, init 3.5) aggregates MHC features. The 64-dimensional TCR and MHC embeddings and their element-wise product form a 192-dimensional fusion vector, processed through a 256-unit dense layer (ReLU, dropout 0.3) and sigmoid output.

### Training procedure

The model was trained with asymmetric focal loss (γ_pos_ = 0.05, γ_neg_ = 1.5) on soft labels (0.25 within-gene negatives, 0.0 cross-gene). Positives were weighted at 1.6× and negatives at 0.8×, with class-balanced allele upweighting (β = 0.999, cap 3.0×). The objective included a PositivePull regularization (λ = 0.05). The model was trained with the Adam optimizer (lr 10⁻³, gradient clip 1.0, batch 256), up to 60 epochs with early stopping (patience 8) on validation AUC. LR reduced 0.5× on plateau (patience 4, min 10⁻⁵). Six-fold cross validation was run on 90% of data; all folds share the same 10% held-out set.

### Evaluation metrics

AUC was computed from ensemble predictions (mean across six folds), stratified by allele, chain status, and gene group. 95% confidence intervals were estimated by bootstrapping (2,000 iterations). Generalization was assessed on the IMMREP 2023 benchmark, restricted to TCRs absent from all training folds and the held-out set.

### Comparison with DePTH2.0

DePTH2.0^13^ was run with published pretrained weights and applied on TCRβ sequences (≤27 aa, standard residues only) from the held-out set and IMMREP benchmark, using separate class I and class II models. TRIOPS was evaluated on the identical sequences using TCRβ-only and paired chain inputs. DePTH2.0 provides no paired-chain mode, so paired-input results report additional TRIOPS capability rather than a matched comparison. AUC was computed identically for both methods, including positive/negative definition, allele stratification, and 2,000-iteration bootstrap CIs, reported separately for class I and class II.

### Unseen-HLA generalization

For this analysis the data were repartitioned separately from the main split, grouping on the MHC pseudosequence (the sole input to the MHC branch) rather than on individual pairs. Whole pseudosequence groups were assigned at random to a fixed held-out set comprising ∼10% of records; groups each exceeding 10% of the data were retained in training to preserve signal.The held-out set comprised 56,703 pairs (9.3% binders) spanning 15 pseudosequences and 16 HLA alleles. The remaining data was divided into six cross-validation folds (k = 6, seed 42), used for early stopping only, so inner-fold validation could share alleles with training while the external held-out set remained pseudosequence-disjoint. Architecture, encoding, masking, loss, and optimization matched the primary model. Held-out pairs were scored by the six-fold ensemble; ROC AUC was computed on pooled pairs and per allele. The unweighted macro-average AUC across alleles is reported as the allele-balanced summary.

#### HLA loci-specific false positive score calibration

For each of the six HLA gene loci (A, B, C, DR, DP, DQ), an ROC curve was constructed from the validation predictions of each fold, and the threshold yielding a 20% false-positive rate was found by linear interpolation. We chose this operating point to equalize the per-locus false-positive rate rather than sensitivity. Because the six gene groups occupy different score scales, their raw thresholds are not directly comparable; subtracting each locus’s 20% FPR threshold rescales every score as a margin above a shared error rate, making the rectified margins comparable across loci. The locus-specific threshold was the mean of the six per-fold thresholds. For each clonotype, each fold score was offset and rectified as max(0, raw − threshold) and the six margins were then averaged, with the gene group inferred from the allele. The per-gene-group threshold is a reference operating point, not a false-positive guarantee on patient repertoires; subtracting it aligns class I and class II at a common operating point, after which the engaged allele is selected among those a patient carries.

#### Embedding visualization and coherence analysis

Fold-4 fusion embeddings were extracted for positive validation examples (binder score ≥ 0.5) from the joint TCR–MHC concatenation layer. Embeddings were L2-normalized and projected to two dimensions with t-SNE using a cosine metric, perplexity of 30, and PCA initialization. Embedding coherence was quantified with four metrics computed on alleles represented by at least 20 examples: kNN purity (k = 15), the adjusted Rand index (ARI) and normalized mutual information (NMI) of a 30-cluster k-means partition relative to allele identity, and the cosine silhouette score. Each metric was compared against a null distribution generated by 1,000 random permutations of allele labels. Separation ratios were defined as the mean between-group cosine distance divided by the mean within-group cosine distance, computed at both the allele and MHC-class levels, with 95% confidence intervals obtained by bootstrap resampling.

#### Positional saliency analysis

Input-embedding gradients were computed from the fold-4 model as the magnitude of the gradient of the predicted binding score with respect to the input embeddings, averaged across embedding dimensions and across examples. TCR saliency was computed per sequence position and stratified by MHC class (class I and class II) and by chain status (paired TCRαβ, TCRβ only, and TCRα only). MHC saliency was computed per pseudosequence position for three representative alleles, HLA-A*02:01, HLA-B*08:01, and HLA-DPA1*01:03/DPB1*04:01, with saliency profiles normalized to their per-allele maximum.

#### TCGA pan-cancer analysis

TCGA bulk RNA-seq was obtained from the Genomic Data Commons^29^. TCGA TCRβ repertoire reconstruction data was obtained from Dr. Li Song. All TCRβ clonotypes were paired using TRIOPS with each patient’s HLA alleles, and six-fold ensemble predictions were calibrated by locus-specific FPR thresholds, with each clonotype assigned to its argmax allele. Immune cell abundances were estimated from LUAD RNAseq data with CIBERSORT^33^ using the LM22 signature matrix, applied to LUAD RNA-seq quantified as transcripts per million (TPM). TCR-HLA engagement was correlated against total T cells (CD4+ plus CD8+) by Pearson r, both unweighted and read-count-weighted; Spearman correlations yielded similar results. Clonal dominance was quantified as Simpson’s D = Σpᵢ².

#### Statistical analysis

Pearson correlations between total T-cell proportion and predicted class I or class II clonotype counts were computed with SciPy, with 95% confidence intervals obtained by bootstrap resampling over 10,000 iterations. Group comparisons were assessed with the Wilcoxon rank-sum test, and effect sizes were quantified with Cliff’s δ. The common-language effect size (CLES) was reported as the probability that a randomly chosen class I-targeting sample shows greater dominance than a randomly chosen class II-targeting sample.

#### Engagement and allele-specific loss

We applied the engagement–loss analysis to the same 8,906-sample TCGA pan-cancer TCRβ repertoire used above (Fig. 4), pairing each patient’s TRIOPS predictions with allele-resolved RNA-seq B-allele frequency (BAF). TCR engagement at each allele was defined as the summed TCRβ read count of the clonotypes assigned to that allele, where each clonotype in a patient was assigned to the single allele with the highest calibrated TRIOPS binding margin across that patient’s genotype (cross-locus argmax). The continuous loss measure was loss = 1 − 2e, where e is the expressed fraction (BAF) of the more-engaged allele, so that loss > 0 marks under-expression of the engaged allele and loss < 0 marks its retention. Informative events were those with |loss| > 0.15 (equivalently BAF outside 0.425 to 0.575), a threshold set to exclude near-symmetric BAF noise.

#### Heterozygous locus selection and quality control

Because allele-specific loss is only defined when a locus presents two distinguishable alleles, the analysis was restricted to heterozygous loci: the class I genes HLA-A, HLA-B, and HLA-C, and the class II β-chain genes HLA-DRB1, HLA-DQB1, and HLA-DPB1. Homozygous loci, which present a single allele and so admit no lost-versus-retained contrast, were excluded. For class II, heterozygosity and BAF were assessed on the β-chain gene of each dimer, the polymorphic chain that defines the expressed allele pair. A heterozygous allele pair was retained when at least one allele carried nonzero engagement and the pair passed all quality-control filters: at least 100 combined RNA-seq reads spanning the locus, a patient repertoire of at least 10 unique TCRβ clonotypes and at least 30 total TCRβ reads, and a cancer type with at least 20 evaluable patients. After filtering, 5,233 patients contributed engagement assignments, yielding 24,936 informative heterozygous pairs across 23 cancer types. All models adjusted for ABSOLUTE tumor purity and log₁₀ total RNA-seq read depth (per cancer distributions, Supplementary Fig 7), mean-centered within each fit and retained only where finite with nonzero variance.

#### Modeling TCR-HLA engagement and concordance

For each cancer type we fit a Jeffreys-penalized (Firth) logistic regression of engaged-allele loss on the two centered covariates. Because both covariates were centered within each cancer, the per-cancer concordance cⱼ = invlogit(intercept) is the purity- and depth-adjusted probability, evaluated at that cancer’s mean purity and depth, that the engaged allele is the lost one; cⱼ > 0.5 denotes preferential loss of engaged alleles and cⱼ < 0.5 preferential retention. A cancer contributed cⱼ only with at least 15 informative events. The pan-cancer estimate aggregates each cancer once, c = invlogit(mean of the per-cancer intercepts), with a 95% confidence interval from a t-distribution on the logit scale (J − 1 degrees of freedom, J contributing cancers) back-transformed through the inverse logit. A two-sided exact binomial sign test of the number of cancers with cⱼ > 0.5 against an expected 0.5 provides a reproducibility test across cancers, independent of the per-cell tests. Per cancer-by-locus significance is the penalized likelihood-ratio test of the intercept (null cⱼ = 0.5), with Benjamini-Hochberg control at q < 0.05 across all finite per-cancer-by-locus cells; asterisks mark class I cells passing this threshold.

#### Within-tumor paired analysis

For each class I locus and cancer type we compared TCRβ engagement at the lost versus retained allele of these informative pairs, with the lost allele defined directly as the under-expressed member (BAF < 0.5), independent of engagement. Per-allele engagement was winsorized at the 95th percentile, log(1 + x) transformed, averaged within each patient, and summarized as the per-cancer median across patients. The cross-cancer comparison was a two-sided exact binomial sign test of the number of cancer types with higher median engagement at the lost allele, among those with a nonzero median difference.

#### Software and computational environment

All models were implemented in Python 3.12.13 using TensorFlow 2.20.0 with Keras 3.13.2. Sequence distance calculations used the python-Levenshtein library. Data processing and downstream analysis used NumPy 2.0.2, pandas 2.2.2, scikit-learn 1.6.1, and SciPy 1.16.3; figures were generated with Matplotlib 3.10.0 and seaborn 0.13.2. Training and inference were performed on an NVIDIA L4 GPU (24 GB, compute capability 8.9) in Google Colab, with TensorFlow built against CUDA 12.5.1 and cuDNN 9.

#### Data availability

Experimentally validated TCR–pMHC binding data were obtained from publicly available databases: TCR3d (https://tcr3d.ibbr.umd.edu), VDJdb (https://vdjdb.cdr3.net), IEDB (https://www.iedb.org), McPAS-TCR (http://friedmanlab.weizmann.ac.il/McPAS-TCR/), and PATCRdb (https://patcrdb.org). The IMMREP 2023 benchmark dataset is available at https://github.com/viragbioinfo/IMMREP_2023_TCRSpecificity. TCGA-LUAD bulk RNA-seq data are available through the NCI Genomic Data Commons (https://portal.gdc.cancer.gov). HLA genotypes were obtained from the TCGA pan-cancer consensus HLA typing benchmark dataset^30^ (https://doi.org/10.1002/1878-0261.12895).

#### Code Availability

TRIOPS is available for non-commercial academic use. An inference pipeline and trained model weights will be released upon publication under an academic-use license at https://github.com/jonssonlab/triops. Use for commercial purposes requires a separate license. Code is available to reviewers during peer review.

### Author contributions

NRR and VDJ conceived the study. NRR curated data for the study, developed the methodology, wrote the software and conducted formal analysis. CMR performed TCGA HLA genotype calls and pan-cancer allele-specific expression quantification. LM performed deconvolution analysis and statistical analyses. VDJ supervised the work. All authors provided feedback on manuscript and figures.

## Acknowledgments

We thank the UC Santa Cruz Genomics Institute for computing resources. This work was supported by a UC Cancer Research Coordinating Committee grant and the Hellman Fellows grant to V.D.J. We thank Li Song for providing the TCGA TCR data and Joshua Stuart for valuable feedback on the manuscript.

---

**Supplementary Figure 1.**
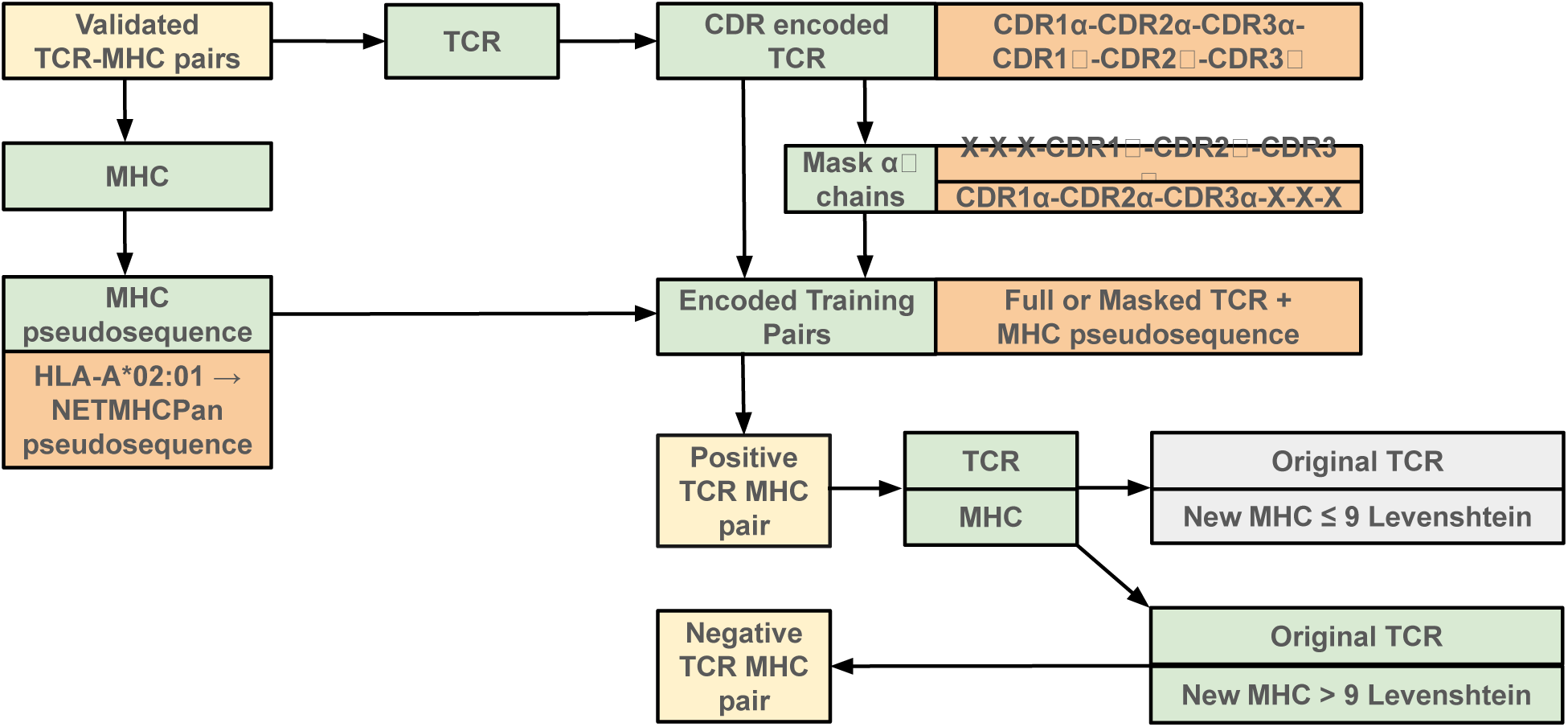
Training data generation pipeline. Schematic of CDR encoding, chain masking, MHC pseudosequence representation, and cross-reactivity–aware negative sampling. Pairs at Levenshtein distance ≤9 are rejected; distance ≥10 retained.

**Supplementary Figure 2.**
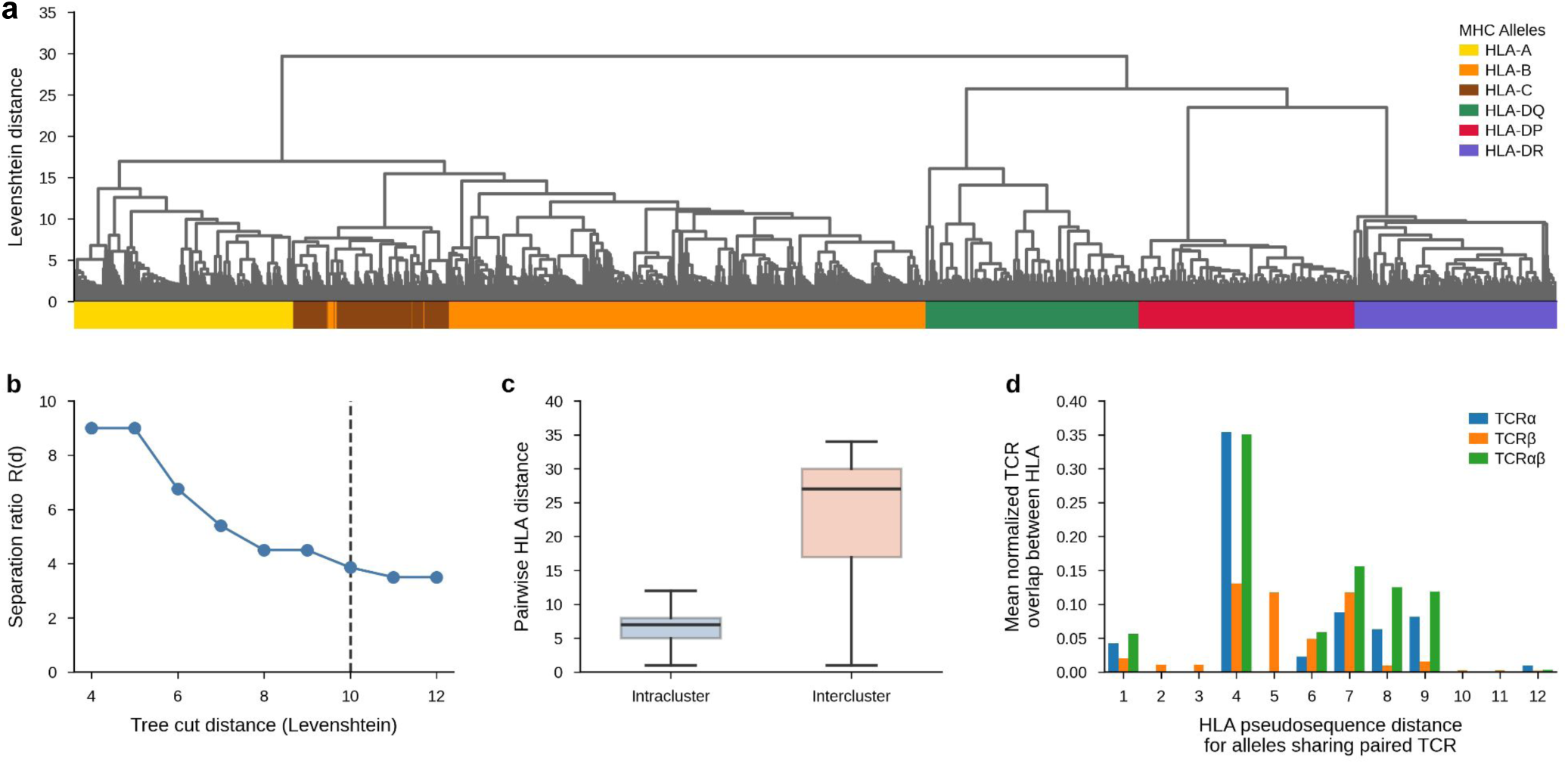
HLA pseudosequence similarity and shared TCR recognition. a, Hierarchical clustering of MHC alleles by Levenshtein distance of NetMHCpan pseudosequences (average linkage). Leaves colored by gene group. b, Cluster separation ratio R(d); plateau at d = 10 (dashed lines). c, Pairwise distances for intracluster and intercluster pairs at cut height d = 10. d, Mean normalized TCR overlap between alleles as a function of pseudosequence distance for TCRα, TCRβ, and paired TCRαβ. Monotonic decrease validates the cross-reactivity–aware sampling threshold.

**Supplementary Figure 3.**
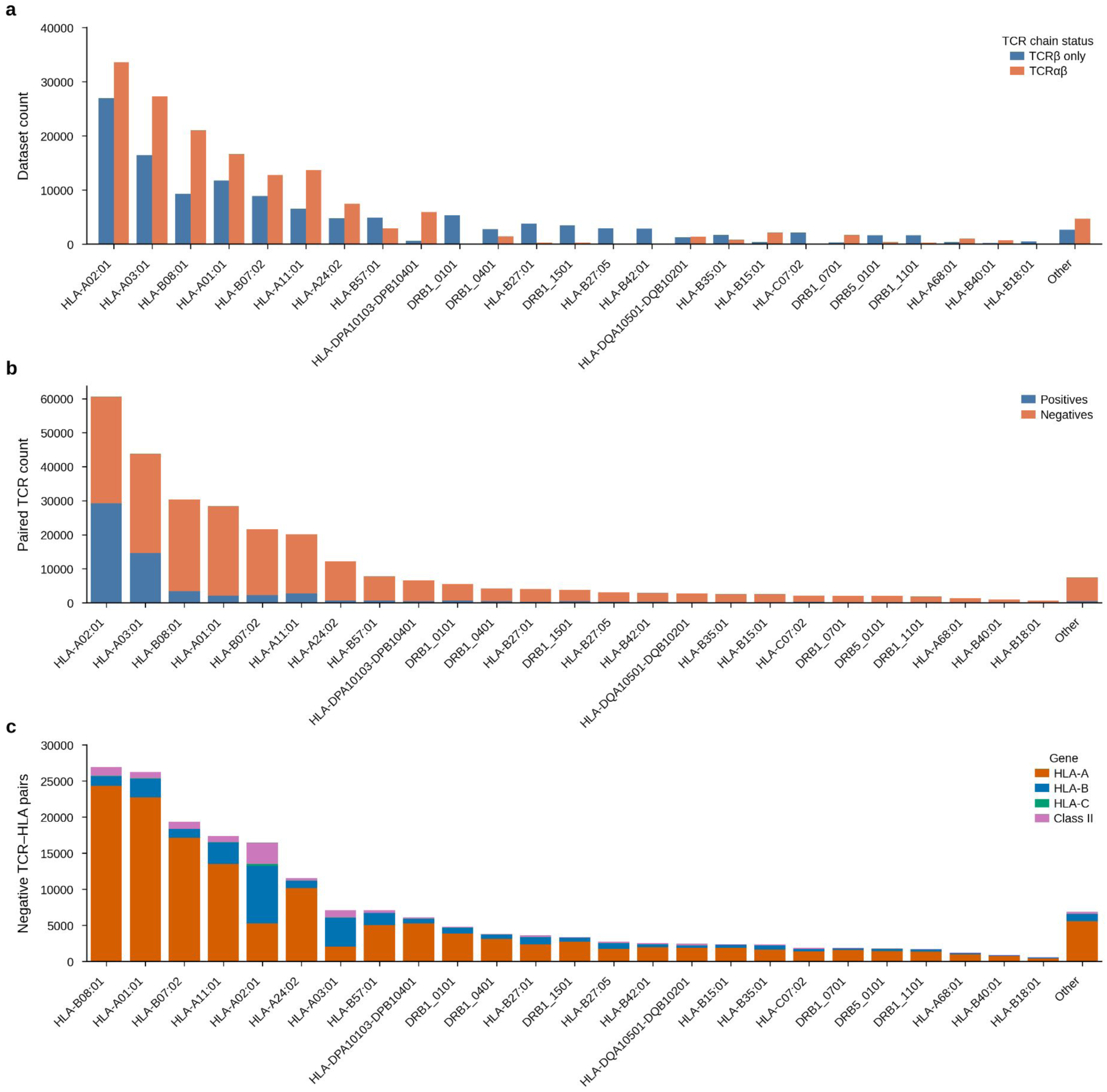
TRIOPS training dataset composition. a, TCR–HLA pair counts by allele, stratified by chain status (TCRαβ vs. TCRβ-only). b, Positive and negative pair counts per allele. c, Negative pair distribution by original HLA gene group, colored by assigned negative allele gene group.

**Supplementary Figure 4.**
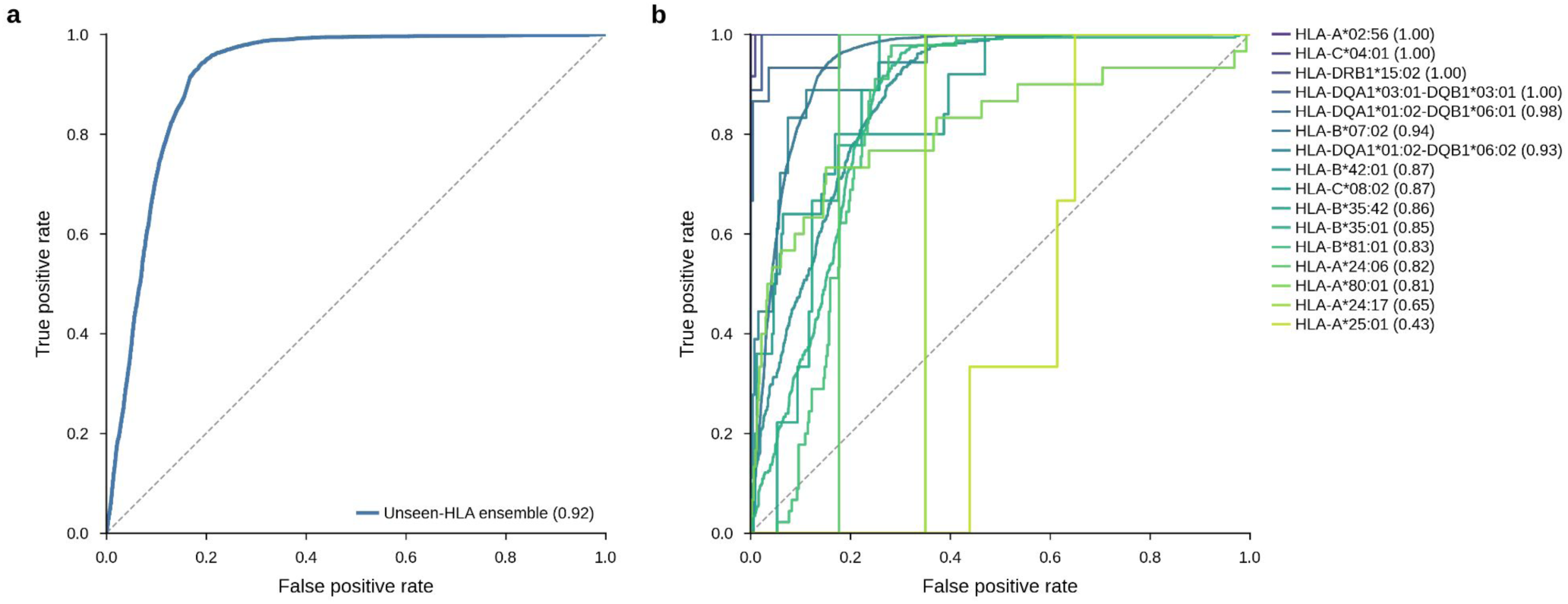
Generalization to HLA alleles unseen during training. 15 pseudosequences (16 HLA alleles) were withheld from training across six cross-validation folds (n = 56,703 TCR–MHC pairs, 9.3% binders). TCRs were not held out, isolating MHC novelty. a, Receiver operating characteristic (ROC) curve for the six-fold ensemble on the holdout (AUC = 0.92; dashed line, chance). b, Per-allele ROC for the 16 held-out alleles, ordered by descending AUC. Per-allele AUC ranged from 0.43 (HLA-A*25:01) to 1.00 (HLA-A*02:56 and HLA-C*04:01); macro-average AUC = 0.87. HLA-B*07:02 accounted for 83% of pairs, so the macro-average provides the allele-balanced summary. Line color denotes allele and follows the AUC ordering.

**Supplementary Figure 5.**
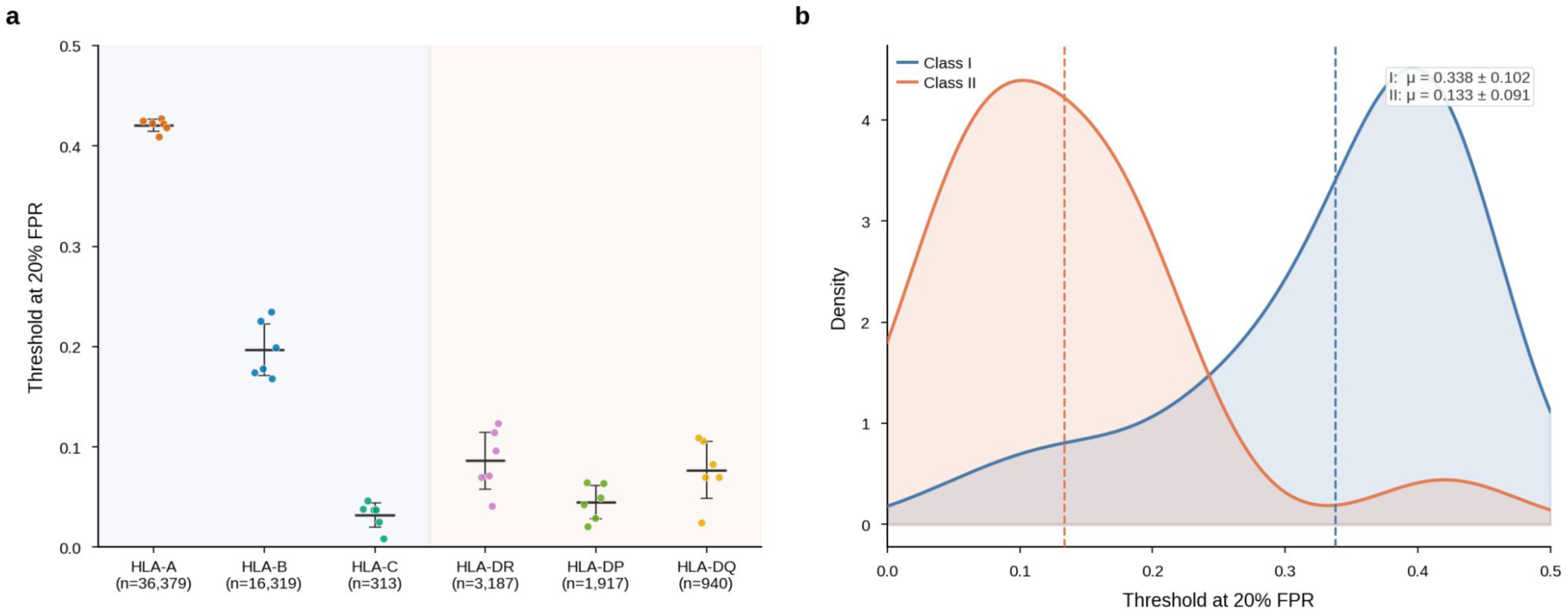
FPR calibration stability across folds. a, Per-locus 20% FPR thresholds across six cross-validation folds for all HLA gene groups (per-locus sample sizes on the x-axis). Points, individual folds; bars, mean ± s.d. Low inter-fold variance indicates robust calibration. b, Threshold distributions by MHC class. Class I, μ = 0.338 ± 0.102; class II, μ = 0.133 ± 0.091 (dashed lines, class means).

**Supplementary Figure 6.**
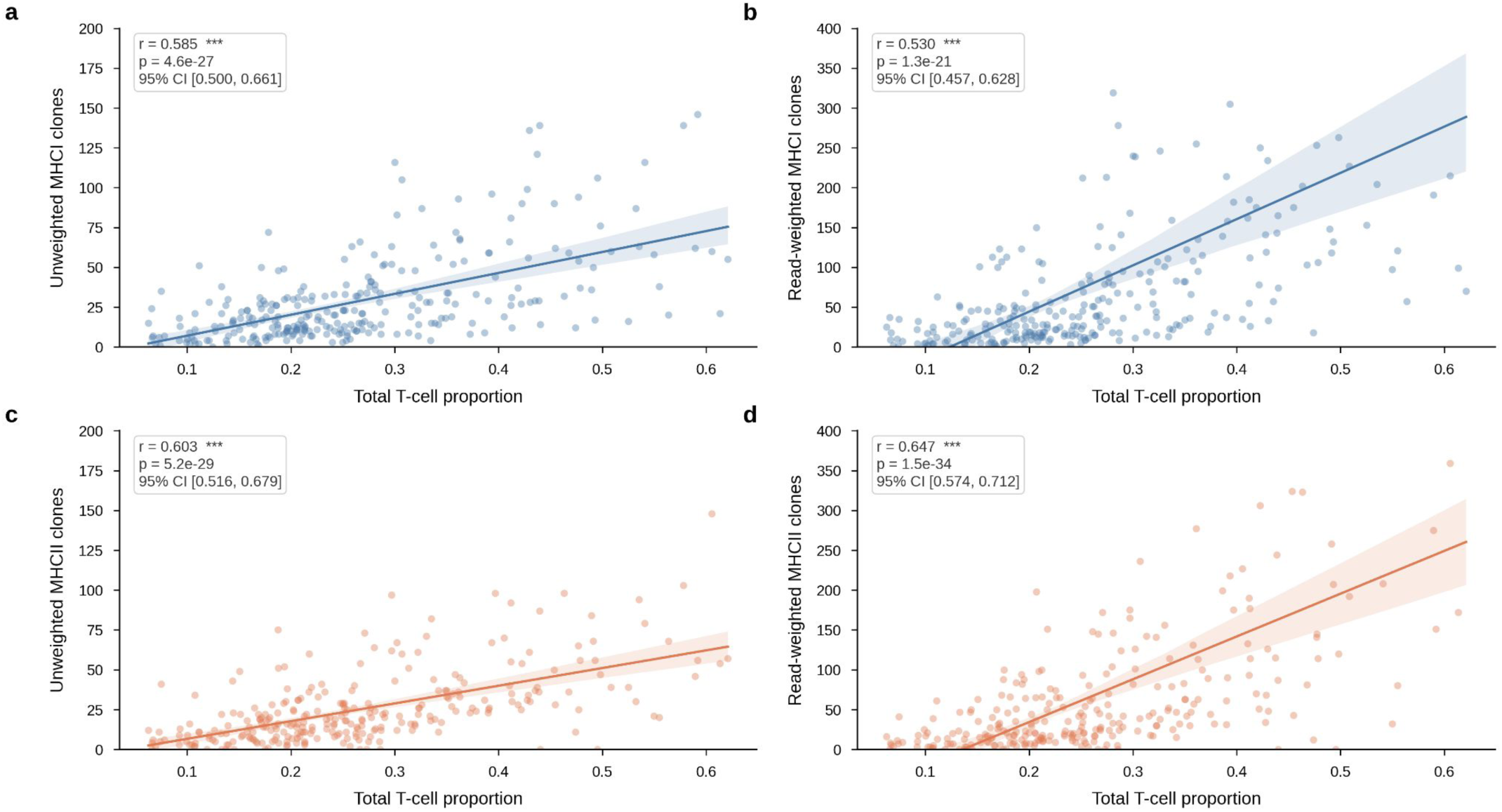
TRIOPS-predicted MHC-targeting clone counts correlate with CIBERSORT-estimated T cell proportions. a, Unweighted class I clone counts versus total T cell proportion (r = 0.585, P = 4.6 × 10⁻²⁷). b, Read-weighted class I clone counts versus total T cell proportion (r = 0.530, P = 1.3 × 10⁻²¹). c, Unweighted class II clone counts versus total T cell proportion (r = 0.603, P = 5.2 × 10⁻²⁹). d, Read-weighted class II clone counts versus total T cell proportion (r = 0.647, P = 1.5 × 10⁻³⁴). Read-count weighting strengthens the class II correlation (c versus d), but attenuates the class I correlation (a versus b). Shaded regions, 95% confidence intervals; n = 275 TCGA-LUAD patients with available CIBERSORT estimates.

**Supplementary Figure 7.**
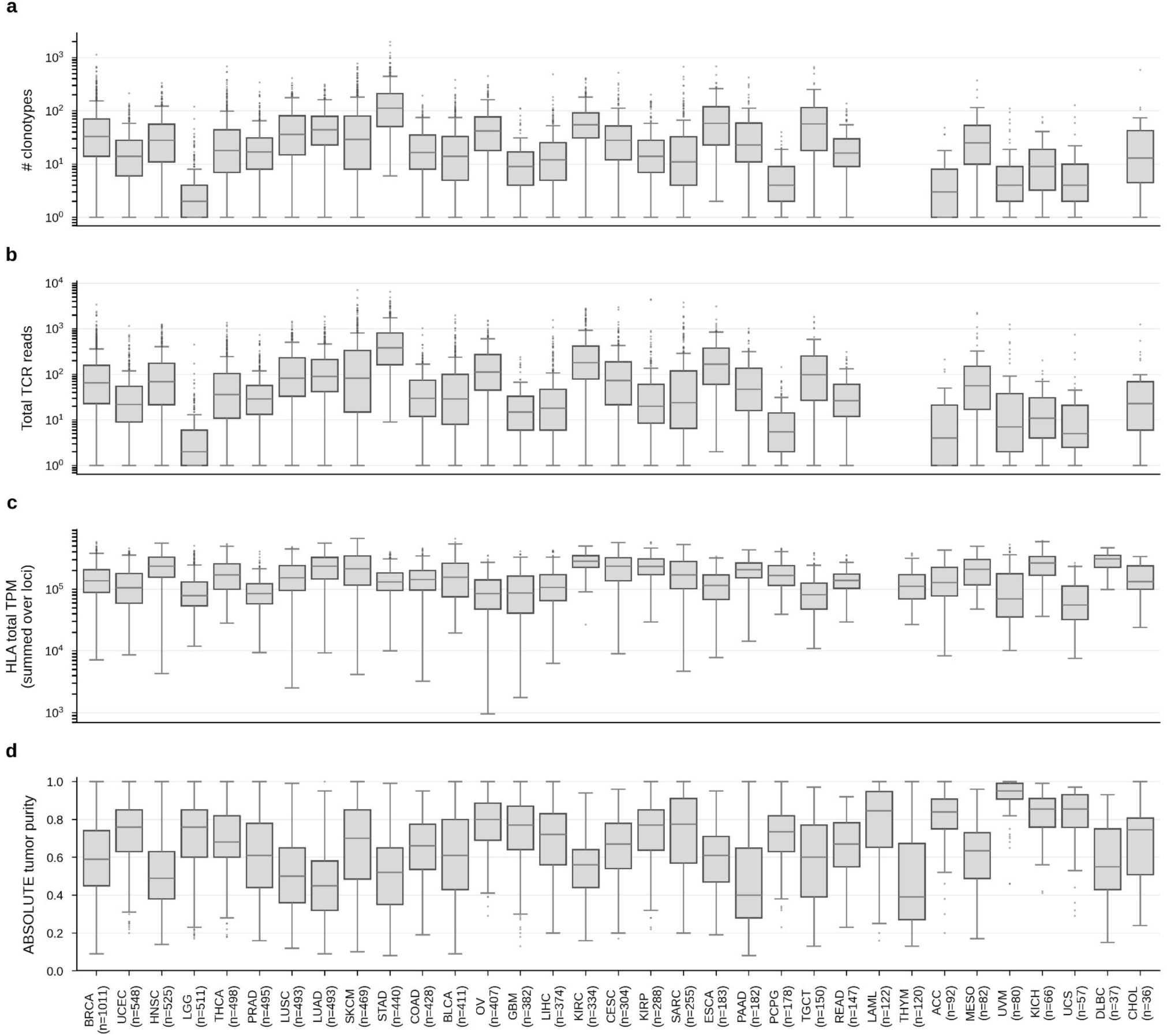
Pre-filter distributions of immune and tumor covariates across TCGA cancer types. Per-cancer distributions of the quantities entering the engagement–loss analysis, shown before quality-control filtering (no minimum TCRβ read, clonotype, locus-read, or per-cancer patient thresholds). Each box is one cancer type (sample sizes in parentheses on the x-axis), showing the median and interquartile range with whiskers at 1.5× IQR and outliers as points; cancers are ordered by sample count. a, Number of recovered TCRβ clonotypes per sample (log scale). b, Total TCRβ read count per sample (log scale). c, Total HLA expression per sample, summed across loci (TPM, log scale). d, ABSOLUTE tumor purity per sample (linear scale, tumor samples only).

